# Balancing signal and photoperturbation in multiphoton light-sheet microscopy by optimizing laser pulse frequency

**DOI:** 10.1101/2020.06.02.130377

**Authors:** Vincent Maioli, Antoine Boniface, Pierre Mahou, Júlia Ferrer Ortas, Lamiae Abdeladim, Emmanuel Beaurepaire, Willy Supatto

## Abstract

Improving the imaging speed of multiphoton microscopy is an active research field. Among recent strategies, light-sheet illumination holds distinctive advantages for achieving fast imaging *in vivo*. However, photoperturbation in multiphoton light-sheet microscopy remains poorly investigated. We show here that the heart beat rate of zebrafish embryos is a sensitive probe of linear and nonlinear photoperturbations. By analyzing its behavior with respect to laser power, pulse frequency and wavelength, we derive guidelines to balance signal and photoperturbation. We then demonstrate one order-of-magnitude signal enhancement over previous implementations by optimizing the laser pulse frequency. These results open new opportunities for fast live tissue imaging.

## 1. Introduction

Multiphoton microscopy has demonstrated unique advantages for deep and live tissue imaging [1]. However, the acquisition speed of standard point-scanning two-photon-excited fluorescence (2PEF) microscopy is generally bounded to a μs per pixel due to signal limitations resulting from fluorophore photophysics [2]. This limit sets important constraints for investigating fast biological phenomena or for multiscale imaging [3]. Improving the speed of multiphoton microscopy is therefore an active field of research [4–9]. Among the strategies developed to improve the acquisition speed in multiphoton fluorescence imaging, light-sheet illumination exhibits distinctive advantages [4], with applications in cell biology [10], neurosciences [11], developmental biology [12], or organoid research [13]. Light-sheet illumination involves reduced average power and peak intensity [4] compared to other approaches based on a collinear arrangement of illumination and detection such as multifocal point-scanning [5] or scanless wide field illumination [9]. Indeed, the parallelization of the illumination in an orthogonal geometry such as light-sheet imaging is done along the light propagation direction, requiring a single excitation beam to excite many pixels. In addition, the low-aperture focusing used to generate the light-sheet results in reduced laser peak intensity at the sample for the same detected signal, without compromising optical sectioning nor axial resolution [4, 14]. Finally, the parallelization of the illumination results in long pixel dwell times and therefore in higher signal levels compared to fast scanning approaches [8]. These characteristics are expected to set different photoperturbation-related constraints in multiphoton light-sheet microscopy compared to fast collinear geometry approaches.

When optimizing a live fluorescence assay, it is essential to characterize the mechanisms and thresholds of photoperturbations at stake during imaging [15]. Knowledge of the dependence of perturbations on illumination parameters can then be used to balance the level of signal and of unwanted effect. In linear imaging techniques using continuous wave lasers such as confocal microscopy, signal and perturbations are generally simply proportional to the excitation power. By contrast, in multiphoton imaging relying on pulsed excitation, additional illumination parameters can be adjusted to mitigate photoperturbations. Indeed, when the dominant perturbation process has a different nonlinear order than the signal, the signal-to-photoperturbation ratio can be increased by adjusting the laser duty cycle *τ*/*T* either through the laser pulse frequency (or repetition rate) *f* = 1/*T* [16], or through the pulse duration *τ* [17]. For instance, when the dominating perturbation process has an order higher than two, increasing the laser pulse frequency is an efficient strategy for increasing two-photon excited fluorescence (2PEF) while keeping photoperturbation at a constant level [16]. In standard multiphoton microscopy, various perturbation mechanisms have been investigated such as linear absorption and photothermal effects [5, 18], fluorophore saturation [19], fluorophore photobleaching [20, 21], or nonlinear photochemical phenomena [16, 17, 22, 23].

In multiphoton light-sheet microscopy, adjusting the laser pulse frequency remains an attractive yet unexplored strategy to balance signal and photoperturbation. Indeed, most multiphoton light-sheet microscopes described so far [4, 10–13, 24–29] were implemented using mode-locked Ti:sapphire laser at *f*~80 Mhz, which is the most commonly used laser source in multitphoton microscopy. However, this choice is questionable and may not take full advantage of light-sheet illumination and its orthogonal geometry. A better understanding of the experimental parameters governing linear and nonlinear photoperturbation involved in the specific illumination conditions used in multiphoton light-sheet illumination is therefore necessary to bring this imaging modality to its full potential.

In this study, we devised a systematic experimental workflow to assess the nature and level of photoperturbations induced during imaging of the beating heart in live zebrafish embryos with multiphoton light-sheet microscopy (or 2P-SPIM, two-photon single-plane illumination microscopy). We found that monitoring the instantaneous heart beat rate (HBR) is a sensitive probe of both heating and higher-order-multiphoton induced perturbations. We identified the level and nonlinear order of photoperturbations during live imaging as a function of laser mean power, pulse frequency, and wavelength. This analysis enabled us to derive guidelines for determining which pulse frequency provides the optimal balance between signal, linear and nonlinear photoperturbation. In turn, we achieved high-speed *in vivo* two-photon imaging with an order-of-magnitude signal increase compared to previous reports using light-sheet illumination.

## 2. Results

### 2.1 Two- and three-photon excited fluorescence increases with lower pulse frequency in multiphoton light-sheet microcopy

In multiphoton microscopy, the order *n* of any optical process *OP_n_* dictates how it is linked to the laser illumination intensity *I,* since *OP*_*n*_~*I*^*n*^. It can then be shown that *OP_n_*, depends on the laser pulse frequency *f* (or period *T=1/f*), the pulse duration τ, and the mean power *P_mean_* as follows

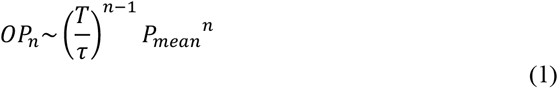

As a consequence, nonlinear signals used in multiphoton light-sheet microscopy, such as 2-photon excited fluorescence (2PEF [4, 24, 27, 28]), second harmonic generation (SHG [30]) or 3-photon excited fluorescence (3PEF [29]) can be enhanced by increasing *T* while keeping *P_mean_* constant. Using 2P-SPIM imaging zebrafish embryos expressing mCherry fluorescent proteins, KTP nanocrystals [30], and fluorospheres embedded in a gel (Supplementary methods and Table S1), we confirmed that both 2PEF and SHG signals depend linearly on *T* (*n* = 2, Fig. 1a-b, Table S2) and that 3PEF signals depend quadratically on *T* (*n* = 3, Fig. 1c). Hence, by decreasing the laser pulse frequency from 80 to 1 MHz at a constant mean power, 2PEF/SHG and 3PEF signals can be increased by a factor 80 and 6400, respectively. Reaching such enhancement factors is however potentially limited by the onset of higher-order unwanted effects caused by pulses of high energy *E_pulse_* and peak power *P_peak_*. In the case of live imaging of zebrafish embryos, we did not observe any saturation of mCherry fluorophore using *P_mean_* = 100 mW and *T* up to 0.2 μs (*f* = 4.4 MHz), corresponding to *E_pulse_* = 28 nJ (Fig. 1a). This value is one-to-two orders of magnitude higher than the saturating *E_pulse_* estimated in point-scanning multiphoton microscopy [19]. Such a difference can be explained by the low illumination focusing used in 2P-SPIM. Indeed, given mCherry two-photon cross section [31] and a 0.1 illumination numerical aperture, the saturating *E_pulse_* should be ~200 nJ according to [19]. While the fluorophore saturation was not limiting, we however observed tissue damage when using higher *T* and *E_pulse_*. As a consequence, it appears that the signal enhancement obtained by tuning the laser pulse frequency is limited by photopertubations, and that optimizing illumination parameters *P_mean_* and *T* requires their systematic investigation.

**Fig. 1.**
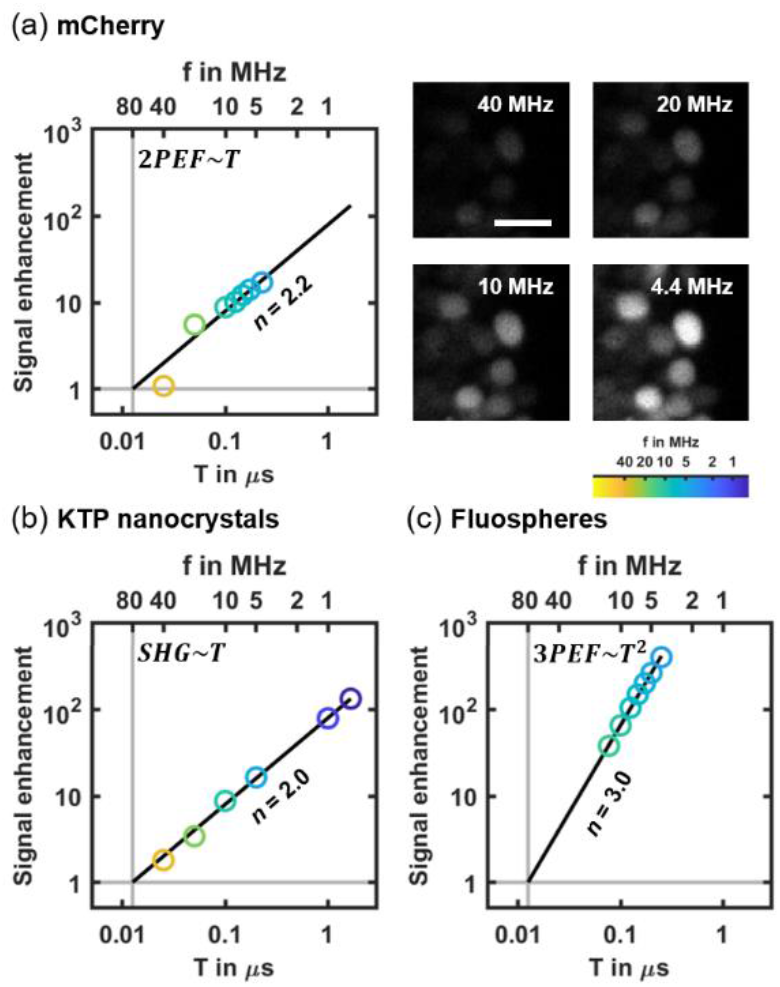
Signal enhancement in multiphoton light-sheet microscopy by decreasing the laser pulse frequency *f* (i.e. by increasing its period *T*) at a given mean power *P_mean_* in the case of 2PEF (a), SHG (b), and 3PEF (c). (a) Linear increase in 2PEF signal (left) from mCherry labeled zebrafish embryos imaged at 168 fps with *P_mean_* = 100 mW (N = 3 embryos). Representative images (right) of mCherry labeled cell nuclei used for signal quantification at *f* = 4.4, 10, 20 and 40 MHz using the same pixel gray scale. Scale bar is 10 μm (b) Linear increase in SHG signals from KTP nanocrystals (N = 500 nanocrystals). (c) Quadratic increase of 3PEF signals from blue fluorescent microspheres (N = 30 microspheres). 2PEF, 3PEF and SHG signals were normalized to the signals at *f* = 80 MHz (vertical gray lines). The order *n* of each optical process is retrieved from a linear fit of logarithmic scaled data (indicated with black lines). Detailed results of linear fits are given in Table S2.

### 2.2 Investigation of linear perturbation and nonlinear photodamage in vivo using zebrafish heart beating as a reporter

To systematically investigate the invasiveness of 2P-SPIM *in vivo* imaging depending on illumination parameters, such as laser pulse frequency *f* or laser mean power *P_mean_*, we designed an experimental workflow using heart dynamic as a reporter of photoperturbation (Fig. 2, Visualization 1 and Supplementary methods). We imaged hearts of zebrafish embryos, using both white-light illumination and 2PEF imaging, during illumination using a femtosecond laser with tunable pulse frequency in the 1-40 MHz range (Fig. 2a). White-light illumination generated images of the heart of sufficient quality to quantify the instantaneous heart beat rate (HBR) even when the femtosecond laser was switched off or when using unlabeled embryos (Fig. 2b-c). We kept all imaging parameters constant, except *P_mean_* and *f = 1/T*. During a typical 130s-experiment (Fig. 2d), the HBR baseline *HBR_0_* was first recorded with the laser switched off. The laser was then turned on during ~40 s and an increase in HBR was observed and quantified when stabilized. Finally, the laser was switched off to follow the HBR recovery (Fig. 2d). We then plotted the relative variation of heart beat rate (*ΔHBR*/*HBR_0_*) as a function of the laser mean power *P_mean_* (Fig. 2e) for different embryos with a fixed pulse frequency *f*. From this graph, we extracted two critical parameters: (i) the linear slope (*S_L_*) characterizing the relative increase in HBR and (ii) the mean power threshold (*P_NL_*) at which we started to observe nonlinear photodamage, such as irreversible arrhythmia, beating arrest or bubble formation (Visualization 1). These two parameters will be used in the next sections to characterize linear and nonlinear invasive effects during 2P-SPIM imaging. We found that this approach was a fast, reproducible and quantitative investigation of optically-induced disruptions. It was also very sensitive and allowed us to detect very small variations of the HBR, down to 0.02 Hz or 1% of relative variation.

**Fig. 2.**
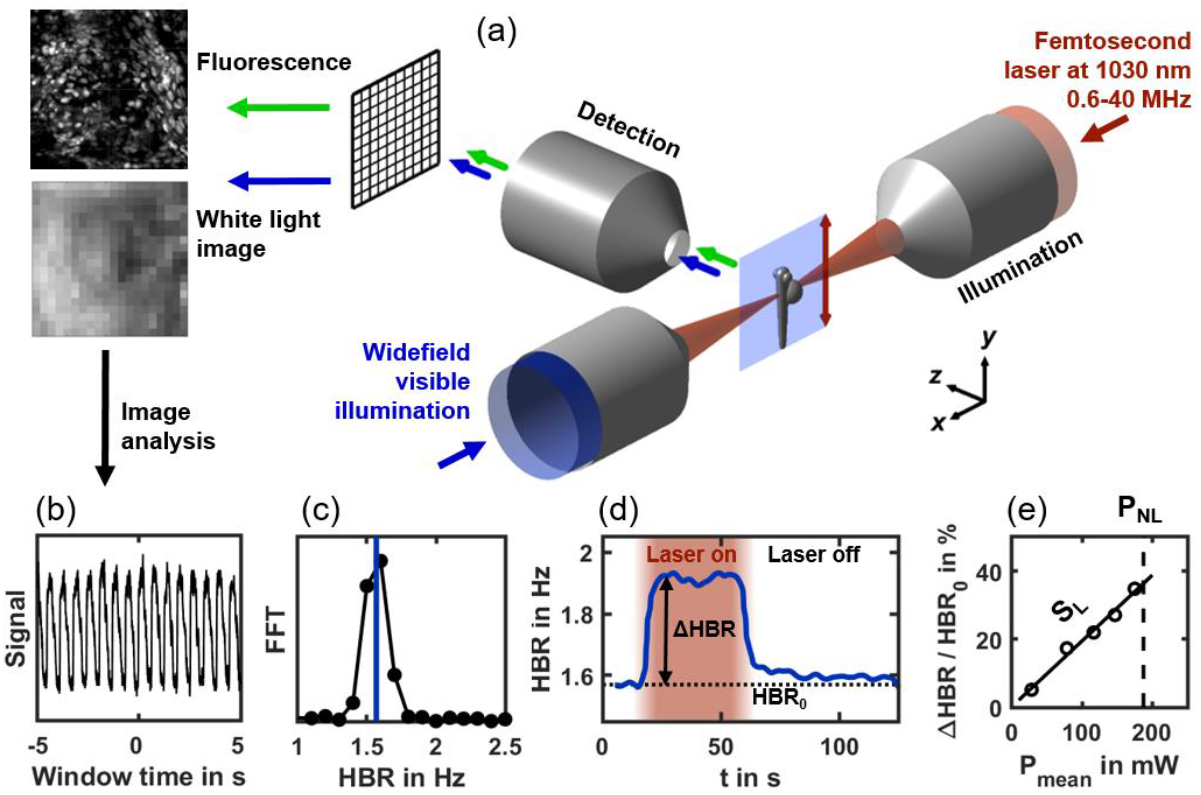
Experimental workflow to quantify linear perturbation and nonlinear photodamage using zebrafish heart beat rate (HBR) as a reporter. (a) 2P-SPIM setup with two illumination modes: a tunable pulse frequency femtosecond laser at 1030 nm wavelength (shown in red) is scanned to generate a light-sheet over the embryo, and wide field white light illumination (shown in blue). Either fluorescence (green arrows) or white light (blue arrows) images are acquired. (b) Periodic intensity fluctuations extracted from white light images correspond to the beating of the heart. (c) Instantaneous HBR is estimated using a ten second windowed Fourier transform of that signal with sub-time point accuracy (blue line). (d) The instantaneous HBR is measured along time, and increases by ΔHBR compared to the baseline HBR_0_ when the fish is exposed to femtosecond laser illumination (in red). (e) At a given laser pulse frequency, the relative ΔHBR is proportional to the laser mean power *P_mean_* with a slope *S_L_*. *P_mean_* is increased up to a power threshold *P_NL_* over which nonlinear photodamage occur estimated from image observation and HBR arrhythmia (Visualization 1).

### 2.3 Linear absorption at low mean power result in limited heating and reversible increase in HBR

When increasing the laser mean power, with all other imaging parameters remaining constant, the first effect we observed was a linear increase in HBR (Fig. 2e). To demonstrate that this effect was due to a reversible linear absorption of the laser light, we conducted several experiments. First, we measured the slope *S_L_* as defined above and showed that it does not depends on the laser pulse frequency (Fig. 3a, and Table S2), which characterized an optical process of order *n* = 1. We verified experimentally that, as expected for a linear effect, *S_L_* does not depends on the position of the focus when moving it away from the center of the heart but keeping the heart within the illumination cone (Fig. 3b). In addition, we observed that this perturbation was reversible: the HBR systematically returned to its base level within a few tens of seconds after illumination stopped (Fig. 2d).

**Fig. 3.**
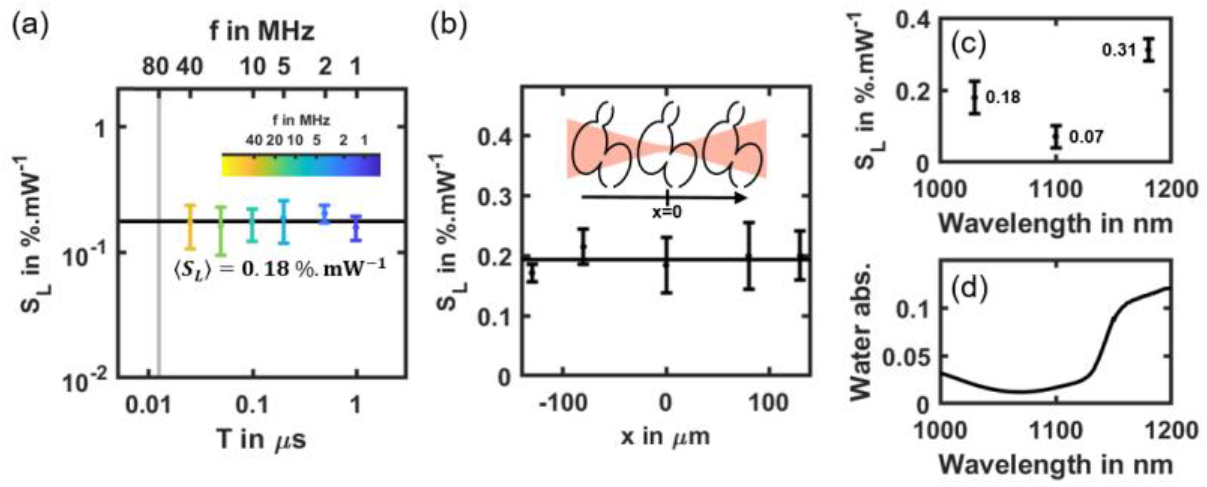
Linear effect at low mean power may be related to water absorption and tissue heating. (a) Linear slope *S_L_* of the HBR relative variation of mCherry labeled zebrafish hearts (N = 27 embryos) depending on the laser pulse frequency *f = 1/T*. Black line indicates mean value <*S_L_*> = 17.7±1.7 %.100 mW^−1^). (b) *S_L_* as a function of the relative position of the fish heart and illumination beam focus in the x-direction at *f* = 10 MHz (N = 3 embryos per position). Black line indicates mean *S_L_* value. (c) *S_L_* at *f* = 80MHz depending laser wavelength. (d) Water absorption spectrum in the 1000-1200 nm wavelength range.

To examine whether this effect was mediated by the fluorophore, we performed the same experiment in unlabeled embryos (Fig. S1a) and in embryos labeled with a different fluorophore (Fig. S1d). We measured similar *S_L_* values in both cases, validating that HBR increase is not mediated by labeling. Finally, to test whether this effect was mediated by water absorption, we repeated the experiments at different wavelengths using a tunable 80MHz laser source. We found that *S_L_* does depend on laser wavelength and follows the same variation as water absorption in the 1000-1200nm range (Fig. 3d). This result suggests that water absorption might be involved and could induce a local heating of the specimen when the laser is on. Since the HBR is known to linearly depend on the temperature in the zebrafish embryo [32, 33], *S_L_* can be related to a change in local temperature. We measured an HBR variation of ~0.29 Hz/°C (Fig. S2) and *S_L_* remains close to 20% per 100 mW when illuminating embryos with *f* = 1-40 MHz and *λ* = 1030nm. As a consequence, we obtain a typical temperature increase of 1.4°C per 100 mW. Such measurement is consistent with recently reported heating measurements in the mouse brain using point-scanning multiphoton microscopy [5, 18]. Together, these results demonstrate that the first detectable disruption when imaging the zebrafish heart with 2P-SPIM is reversible and due to linear absorption of the laser light. They also suggest that this linear effect is not mediated by the fluorophore and could be due to water absorption. Using an illumination of *P_mean_* = 70 mW at λ = 1030 nm would then result in a typical temperature increase of ~1°C and a limited and reversible HBR increase of ~0.3 Hz, which remains within physiological conditions for zebrafish embryos. More generally, it appears that adjusting the laser pulse frequency at a constant mean power is an effective strategy to enhance the signal while keeping linear heating at a constant level.

### 2.4 Scaling law of highly nonlinear photodamage

When increasing further the laser mean power, with all other imaging parameters being held constant, we observed strong and irreversible tissue photodamage at a given power threshold *P_NL_* as defined above (Fig. 2e). To demonstrate that this effect was due to a nonlinear photodamage mechanism, we measured *P_NL_* at different laser pulse frequencies. Unlike in the case of *S_L_* and linear heating due to water absorption, *P_NL_* strongly depends on *T* (Fig. 4a), and decreases from 320 mW at 20 MHz to 30 mW at 1 MHz (Table S1). This dependence on *T* is expected from a nonlinear optical process according to Equation (1). To estimate the scaling law and the order *n* of this perturbation mechanism, we performed a linear fit on *P_NL_(T)* plotted in logarithmic scale (Fig. 4a). According to Equation (1), the threshold *P_NL_* should scale as 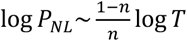. We found *n* = 5.8 (Table S2 for details) with a good reproducibility of the experiment from one embryo to the other considering the high nonlinearity of the optical effect. To confirm the nonlinear nature of these perturbations, we measured *P_NL_* at *f* = 10 MHz for different position of the heart relative to the beam focus along the illumination axis and keeping the heart within the illumination cone. Unlike in the case of *S_L_* and a linear effect, we observed a lower photodamage threshold *P_NL_* when the laser beam was focused at the center of the heart (Fig. 4b). Its variation could be fitted to the intensity profile along the illumination axis of a Gaussian beam of 0.05 numerical aperture (black line in Fig 4b), which is close to the actual numerical aperture of the 2P-SPIM setup. Such behavior is expected from a nonlinear optical process that is sensitive to the local intensity. To test whether the observed nonlinear photodamage were mediated by the fluorophore, we measured *P_NL_(T)* and estimated *n* in unlabeled embryos (Fig. S1a-b) and in embryos labeled with a different fluorophore (Fig. S1c-d). We found similar values for *P_NL_* and the order *n*, which was close to 5 (*n* = 4.8 and 4.9, respectively; Table S2). If photodamage had been sensitive to the fluorophore, we would have obtained significantly higher values of *P_NL_* and *n* in the case of unlabeled embryos. Together, these results demonstrate that increasing the signal in 2P-SPIM by increasing the average power or decreasing the laser is limited by irreversible highly nonlinear photodamage with an order close to *n* = 5, which are not mediated by the fluorophore.

**Fig. 4.**
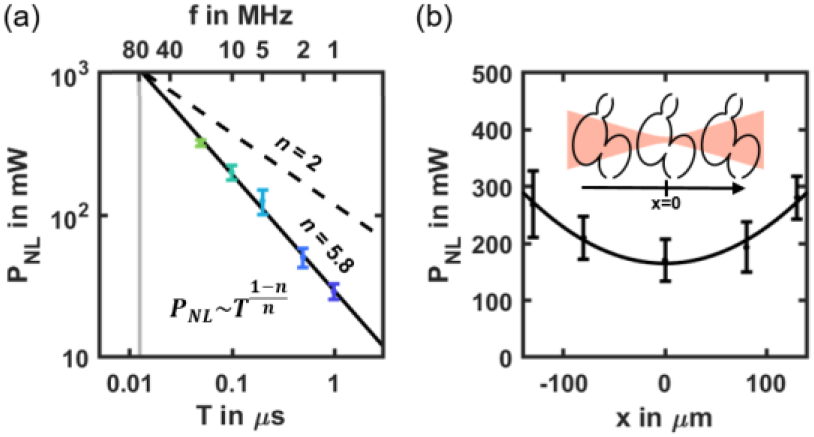
Scaling law of nonlinear photodamage. (a) Nonlinear photodamage threshold *P_NL_* in mCherry labeled zebrafish hearts (N = 21 embryos) depending on the laser pulse frequency *f* = 1/*T*. Black line shows the result of the scaling law fitted on logarithmic scaled data. The *P_NL_* (T) follows a scaling law of order *n*~5.8 (see Table S2 for details). Black dashed line indicates a scaling low of order *n* = 2 to show how it deviates from 2PEF signal. (b) *P_NL_* as a function of the relative position of the heart and illumination beam focus at *f* = 10 MHz (N = 3 embryos per position). Black line indicates the result of a Gaussian fit. Error bars indicate standard deviation.

### 2.5 Photobleaching of mCherry at 1030 nm using 2PSPIM is a nonlinear process

The last unwanted photoinduced effect we analyzed in this study is fluorophore photobleaching. We measured the photobleaching rate of mCherry at 1030 nm wavelength when tuning the laser pulse frequency (Fig. 5). To be relevant to experimental applications, this measurement was performed at a constant starting 2PEF signal level by adjusting the laser power *P_mean_* to compensate for the variation in *T* (see Table S2 for details). It results in 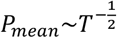 according to (1). At *f* = 40 MHz (yellow line in Fig. 5a), very low photobleaching is observed, which confirms previously reported experiments at *f* = 80 MHz [34]. However, we showed the photobleaching rate increases with T. Since 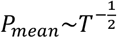, photobleaching of order *n* should scale as 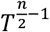. Hence, we estimated the order of the mCherry photobleaching rate to be *n* = 3.3 (Table S2), which is consistent with previous investigations of photobleaching in multiphoton microscopy [20]. In the following section, we will consider the photobleaching observed at *f*=0.6 MHz (dark blue line in Fig. 5a) as the experimental threshold. It corresponds to a 50% signal decay after ~150 images or ~1s of acquisition.

**Fig. 5.**
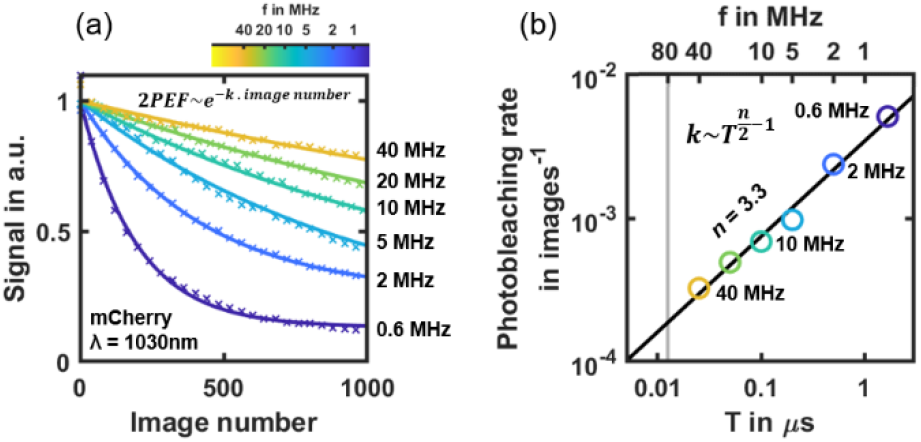
Scaling law of mCherry photobleaching at 1030nm wavelength using 2P-SPIM. (a) mCherry 2PEF signal decay during 2P-SPIM imaging depending on the number of acquired images at different laser pulse frequency f (from 0.6 to 40 MHz). Mean power is adjusted depending on *f* to start with the same 2PEF signal level at each experiment (N = 3 embryos per condition). Solid lines indicate exponential fits used to estimate the photobleaching rate. (b) Photobleaching rate depending on laser pulse frequency *f*. These logarithmic scaled data are used to perform a linear fit (black line) and estimate the *n*-order of the photobleaching rate.

### 2.6 Balancing signal and photoperturbation to optimize live 2P-SPIM imaging of the zebrafish heart

To select the optimal laser pulse frequency and enhance signal during 2P-SPIM imaging, we used our characterization of heating, highly nonlinear photodamage and photobleaching. We represented in a single graph a model of how they limit signal enhancement when adjusting the laser pulse period *T* (or frequency *f* = 1/*T*) and mean power *P_mean_* (Fig. 6a). The signal enhancement is defined as the 2PEF signal normalized to the signal obtained in typical 2P-SPIM imaging conditions (*f_0_* = 1/*T_0_* = 80 MHz and *P_0_* = 70 mW), as previously reported [34]. In this graph, a constant photoperturbation is represented by a straight line whose slope depends on its order. The position of this line and the sign of its slope is critical to optimize the laser pulse frequency and to balance signal and photoperturbation, as explained in Supplementary Methods.

**Fig. 6.**
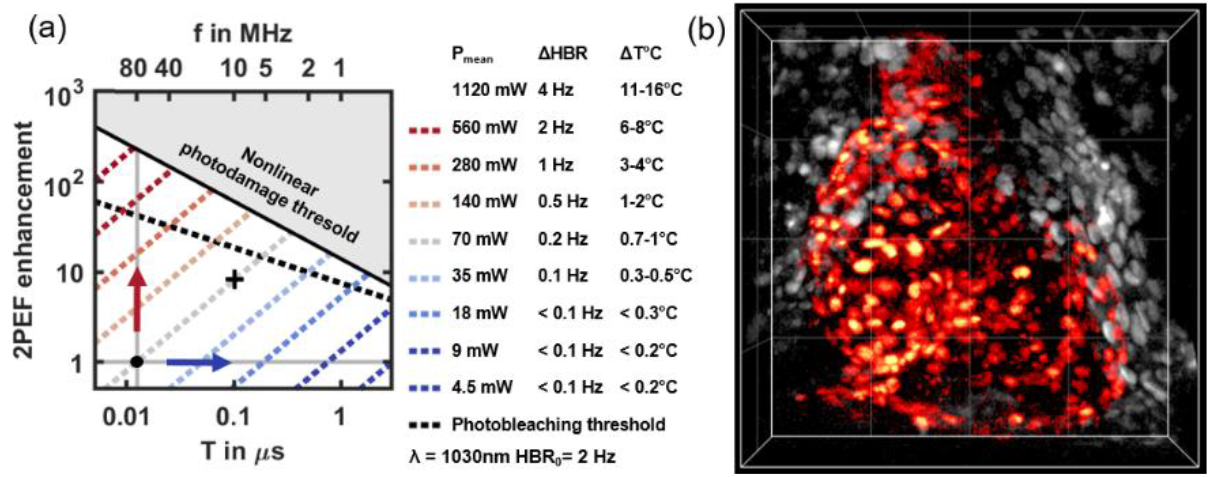
Optimized 2P-SPIM imaging of the zebrafish heart at 10MHz pulse frequency. (a) 2PEF signal enhancement graph used to select the optimal laser pulse frequency. Signal enhancement corresponds to 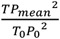 with *T*_0_ = 1/*f*_0_ = 1/80 MHz and *P_0_* = 70 mW, the reference 2P-SPIM imaging conditions (black dot). Solid and dashed black lines correspond to the nonlinear photodamage 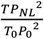 and the photobleaching thresholds, respectively. Dashed blue-to-red lines corresponds to line of constant mean power and constant linear effect (variation of heart beat rate ΔHBR and of temperature ΔT°C). (b) 2P-SPIM imaging of the beating heart in histone mCherry-labeled zebrafish embryo. 4D imaging at 168 fps using *f* = 10MHz and *P_mean_* = 70mW (black cross in a) with 200×200×100μm or 500×500×100 voxels field-of-view and one heart cycle of 75 frames. 4D reconstruction using post-acquisition time synchronization as previousy described [34, 35]. 3-D rendering and manual heart segmentation (red cells) performed with Imaris. Ventral view with anterior up. Grid spacing 50 μm.

Starting from previously reported imaging conditions of 2P-SPIM imaging of the beating heart (black dot in Fig. 6a, at *f_0_* and *P_0_*), several strategies are possible. First, the signal can be increased at *f* = 80 MHz by increasing the mean power, that is following the vertical line on the graph (Fig. 6a, red arrow). In this case, a >200-fold signal enhancement can be obtained before reaching nonlinear photodamage at *P_mean_*>1W. However, large mean powers will induce heating beyond physiological conditions. Our analysis shows that at *f* = 80 MHz, photothermal effects are the limiting factor in 2P-SPIM, which is different to point-scanning multiphoton microscopy [16, 17, 22] and calls for a different optimization approach. It can be explained by the low illumination NA used in 2P-SPIM.

A second possibility would be to reduce *f* down to the 1 MHz range at constant 2PEF signal by following a horizontal line on the graph (Fig. 6a, blue arrow). The graph shows that a constant signal with lower heating can be obtained while reducing the mean power below 10 mW. However, in this regime, the nonlinear photodamage threshold is very low and prevents signal enhancement. In addition, photobleaching at low laser pulse frequencies as reported above for mCherry could be limiting depending on the fluorophore properties.

To optimize heart imaging, we therefore followed an intermediate path. We kept *P_mean_* constant at *P_0_* = 70 mW (light gray dashed line in Fig. 6a). We then decreased *f* and increased the signal while ensuring that heating was less than 1°C therefore keeping the HBR in physiological conditions (less than 0.2 Hz variation). By using *f* = 10 MHz (black cross in Fig. 6a), *P_mean_* remained one order of magnitude below the nonlinear photodamage threshold and we obtained a one order-of-magnitude increase in 2PEF signal. Such signal was sufficient to image the beating heart at the fastest speed of the camera (168 fps, corresponding to 40 MHz pixel rate), without significant photobleaching (Fig 6b, Visualization 2). Indeed, we measured less than 10% signal loss after 75 frames (corresponding here to one heart cycle).

## 3. Discussion

In this study, we established that live imaging of mCherry-labeled zebrafish embryonic heart using 2P-SPIM is significantly optimized by decreasing the laser pulse frequency from 80 to ~10 MHz. This strategy increases the signal by 8-fold without inducing additional heating nor reaching nonlinear photodamage thresholds or significant photobleaching rates. Hence, we achieved high-speed multiphoton imaging *in vivo* while maintaining both low laser average power and peak intensity. Indeed, the mean laser power (*P_mean_* = 70 mW) was at least one order of magnitude lower than used in other parallelized techniques using multifocal point-scanning [5] or wide-field illumination [36], which limits thermal effect. In addition, the laser peak intensity (*I_peak_* = 0.1 TW.cm^2^) was maintained below typical values used in standard multiphoton point-scanning microscopy, and one-to-two orders of magnitude below the values used in recent development of fast multiphoton microscopy [5, 8]. Our study also provides guidelines to further optimize 2P-SPIM imaging. For instance, increasing the laser wavelength from 1030 to the 1100 nm range would drastically reduce linear absorption and heating (Fig. 3c). By using the same reasoning, and assuming that the level and order of nonlinear photodamage are similar, the optimal laser pulse frequency would then be in the range of 20-30 MHz with *P_mean_*~180 mW, resulting in a 2- to 3-fold additional increase of 2PEF signal at constant thermal effects. In addition, mCherry excitation would be enhanced at 1100 nm, with potentially lower photobleaching. However, laser sources with such parameters delivering sufficient mean power are not yet common.

To adjust the laser duty cycle 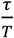 (Eq. 1), one could also reduce the laser pulse width *τ* and potentially obtain similar results than increasing *T*. In this study, we decided to adjust the laser pulse frequency, since using sub-50 fs pulses in multiphoton microscopy brings additional dispersion issues.

This study mainly focused on 2PEF signals. However, a similar signal enhancement strategy can be applied to other nonlinear contrast mechanisms, such as SHG (Fig. 1b) or 3PEF (Fig. 1c and S3), taking into account some subtleties. In the case of multiphoton light sheet imaging of SHG nanocrystals [30], the signal is not limited by saturation or photobleaching and the illumination wavelength can be freely adjusted to minimized water absorption since these probes are achromatic. Additionally, the higher the order of the signal, the stronger its enhancement will be by adjusting the pulse frequency. A quadratic signal enhancement is obtained in the case of 3PEF, corresponding to 64-fold more signal at *f* = 10 MHz (Fig. S3) compared to previously reported 3P-SPIM at *f* = 80MHz [29].

More generally, to take full advantage of light-sheet illumination we demonstrated the laser pulse frequency needs to be optimized compared to current implementations of 2P-SPIM. As a word of caution, it should be noted that some results of this study may be application-specific, such as the photobleaching scaling law of mCherry or the photodamage threshold in zebrafish cardiac tissue. In addition, other experimental parameters may have critical impact on photoperturbations, such as the laser wavelength or the illumination scan speed. However, our study confirmed several general guiding rules. First, unlike in standard point-scanning microscopy, in the case of 2P-SPIM at *f* = 80 MHz, live imaging is not limited by fluorophore saturation or nonlinear photodamage, but by a dominating linear effect, which is likely heating by water absorption. This comes from the lower illumination NA and laser peak intensities used in 2P-SPIM compared to point-scanning techniques. Together, our results suggest *f* = 80 MHz is not adapted to 2P-SPIM imaging and that decreasing *f* to enhance 2PEF signals is a better strategy than simply increasing the laser power. Then, nonlinear photodamage and photobleaching limits how low *f* can be. Finally, laser pulse frequency below 80 MHz and above 1 MHz should be used. Ideally, a laser source providing tunability in both wavelength (900-1200nm) and pulse frequency (typically 4-40 MHz) would be ideal to optimize live 2P-SPIM imaging. The resulting signal enhancement will significantly extend the range of application of live 2P-SPIM imaging, with faster acquisition or stronger signal-to-noise ratio.

## 4. Conclusion

In conclusion, we investigated photoperturbations in live 2P-SPIM imaging by tuning the laser pulse frequency instead of the mean power. By using zebrafish heart beating as a rapid and sensitive experimental reporter, we identified and characterized different photoperturbation processes, specifically a reversible linear effect, likely due to water absorption and heating, and an irreversible nonlinear process of nonlinear order ~5. None of these perturbations were mediated by the fluorophore. In addition, we characterized the scaling law of a fluorophore photobleaching rate. We established that the limiting parameters in 2P-SPIM differ from the ones previously reported using point-scanning multiphoton microscopy [16, 17] and call for a different optimization strategy. Finally, we used these results to balance signal and photoperturbation in live 2P-SPIM imaging of the zebrafish heart: optimizing the laser pulse frequency, we obtained an 8-fold increase in 2PEF signal without inducing additional heating nor reaching nonlinear photodamage thresholds. Such optimization allowed us to image the zebrafish beating heart at the highest camera speed, and with improved signal-to-noise ratio compared to previously reported [34]. Hence, we achieved high-speed multiphoton imaging *in vivo* while maintaining both low laser average power and peak intensity. More generally, these results show that *f* = 80 MHz is not adapted to 2P-SPIM imaging since it causes significant linear absorption and heating. In addition, to enhance 2PEF signals, lowering *f* appears as a better strategy than increasing the laser power. However, reaching the 1 MHz range may expose to nonlinear photodamage and fluorophore photobleaching. In conclusion, it appears that a femtosecond laser source tunable in both wavelength (900-1200 nm) and pulse frequency (4-40 MHz) would be ideal in 2P-SPIM to optimize live imaging conditions. The significant signal improvement obtained by optimizing the laser pulse frequency opens new opportunities for fast *in vivo* imaging with multiphoton light-sheet microscopy.

## Acknowledgments

We thank Marie-Claire Schanne-Klein, Chiara Stringari, and other LOB members for scientific feedbacks, as well as Julien Vermot and his team (IGMC, Illikirch, France and Imperial College London, UK) for scientific discussions. We thank the AMAGEN facility at CNRS Gif-sur-Yvette (UMS 3504 CNRS / UMS 1374 INRA), for providing zebrafish embryos, Emilie Menant and Isabelle Lamarre for zebrafish husbandry, Jean-Marc Sintes and Xavier Solinas for technical support.

## References

1. F. Helmchen and W. Denk, “Deep tissue two-photon microscopy,” Nat Methods 2, 932–940 (2005).

2. J. Mertz, “Molecular photodynamics involved in multi-photon excitation fluorescence microscopy,” European Physical Journal D 3, 53–66 (1998).

3. J. Lecoq, N. Orlova, and B. F. Grewe, “Wide. Fast. Deep: Recent Advances in Multiphoton Microscopy of In Vivo Neuronal Activity,” J Neurosci 39, 9042–9052 (2019).

4. T. V. Truong, W. Supatto, D. S. Koos, J. M. Choi, and S. E. Fraser, “Deep and fast live imaging with two-photon scanned light-sheet microscopy,” Nat. Methods 8, 757–760 (2011).

5. T. Zhang, O. Hernandez, R. Chrapkiewicz, A. Shai, M. J. Wagner, Y. Zhang, C. H. Wu, J. Z. Li, M. Inoue, Y. Gong, B. Ahanonu, H. Zeng, H. Bito, and M. J. Schnitzer, “Kilohertz two-photon brain imaging in awake mice,” Nat Methods 16, 1119–1122 (2019).

6. A. Kazemipour, O. Novak, D. Flickinger, J. S. Marvin, A. S. Abdelfattah, J. King, P. M. Borden, J. J. Kim, S. H. Al-Abdullatif, P. E. Deal, E. W. Miller, E. R. Schreiter, S. Druckmann, K. Svoboda, L. L. Looger, and K. Podgorski, “Kilohertz frame-rate two-photon tomography,” Nat Methods 16, 778–786 (2019).

7. D. R. Beaulieu, I. G. Davison, K. Kilic, T. G. Bifano, and J. Mertz, “Simultaneous multiplane imaging with reverberation two-photon microscopy,” Nat Methods 17, 283–286 (2020).

8. J. Wu, Y. Liang, S. Chen, C. L. Hsu, M. Chavarha, S. W. Evans, D. Shi, M. Z. Lin, K. K. Tsia, and N. Ji, “Kilohertz two-photon fluorescence microscopy imaging of neural activity in vivo,” Nat Methods 17, 287–290 (2020).

9. E. Papagiakoumou, E. Ronzitti, and V. Emiliani, “Scanless two-photon excitation with temporal focusing,” Nat Methods (2020).

10. E. S. Welf, M. K. Driscoll, K. M. Dean, C. Schafer, J. Chu, M. W. Davidson, M. Z. Lin, G. Danuser, and R. Fiolka, “Quantitative Multiscale Cell Imaging in Controlled 3D Microenvironments,” Dev Cell 36, 462–475 (2016).

11. S. Wolf, W. Supatto, G. Debregeas, P. Mahou, S. G. Kruglik, J. M. Sintes, E. Beaurepaire, and R. Candelier, “Whole-brain functional imaging with two-photon light-sheet microscopy,” Nat Methods 12, 379–380 (2015).

12. G. T. Reeves, N. Trisnadi, T. V. Truong, M. Nahmad, S. Katz, and A. Stathopoulos, “Dorsal-Ventral Gene Expression in the Drosophila Embryo Reflects the Dynamics and Precision of the Dorsal Nuclear Gradient,” Dev Cell 22, 544–557 (2012).

13. D. J. Richards, Y. Li, C. M. Kerr, J. Yao, G. C. Beeson, R. C. Coyle, X. Chen, J. Jia, B. Damon, R. Wilson, E. Starr Hazard, G. Hardiman, D. R. Menick, C. C. Beeson, H. Yao, T. Ye, and Y. Mei, “Human cardiac organoids for the modelling of myocardial infarction and drug cardiotoxicity,” Nat Biomed Eng 4, 446–462 (2020).

14. W. Supatto, T. V. Truong, D. Débarre, and E. Beaurepaire, “Advances in multiphoton microscopy for imaging embryos,” Curr. Opin. Genet. Dev. 21, 538–548 (2011).

15. P. P. Laissue, R. A. Alghamdi, P. Tomancak, E. G. Reynaud, and H. Shroff, “Assessing phototoxicity in live fluorescence imaging,” Nature Methods 14, 657–661 (2017).

16. N. Ji, J. C. Magee, and E. Betzig, “High-speed, low-photodamage nonlinear imaging using passive pulse splitters,” Nat Methods 5, 197–202 (2008).

17. D. Debarre, N. Olivier, W. Supatto, and E. Beaurepaire, “Mitigating phototoxicity during multiphoton microscopy of live Drosophila embryos in the 1.0-1.2 microm wavelength range,” PLoS One 9, e104250 (2014).

18. K. Podgorski and G. Ranganathan, “Brain heating induced by near-infrared lasers during multiphoton microscopy,” J Neurophysiol 116, 1012–1023 (2016).

19. K. Charan, B. Li, M. Wang, C. P. Lin, and C. Xu, “Fiber-based tunable repetition rate source for deep tissue two-photon fluorescence microscopy,” Biomed Opt Express 9, 2304–2311 (2018).

20. G. H. Patterson and D. W. Piston, “Photobleaching in two-photon excitation microscopy,” Biophys J 78, 2159–2162 (2000).

21. G. Donnert, C. Eggeling, and S. W. Hell, “Major signal increase in fluorescence microscopy through dark-state relaxation,” Nat Methods 4, 81–86 (2007).

22. A. Hopt and E. Neher, “Highly nonlinear photodamage in two-photon fluorescence microscopy,” Biophys J 80, 2029–2036 (2001).

23. W. Supatto, D. Debarre, B. Moulia, E. Brouzes, J. L. Martin, E. Farge, and E. Beaurepaire, “In vivo modulation of morphogenetic movements in Drosophila embryos with femtosecond laser pulses,” Proc Natl Acad Sci U S A 102, 1047–1052 (2005).

24. J. Palero, S. I. Santos, D. Artigas, and P. Loza-Alvarez, “A simple scanless two-photon fluorescence microscope using selective plane illumination,” Opt Express 18, 8491–8498 (2010).

25. R. Tomer, K. Khairy, F. Amat, and P. J. Keller, “Quantitative high-speed imaging of entire developing embryos with simultaneous multiview light-sheet microscopy,” Nat Methods 9, 755–763 (2012).

26. F. O. Fahrbach, V. Gurchenkov, K. Alessandri, P. Nassoy, and A. Rohrbach, “Light-sheet microscopy in thick media using scanned Bessel beams and two-photon fluorescence excitation,” Opt Express 21, 13824–13839 (2013).

27. F. Cella Zanacchi, Z. Lavagnino, M. Faretta, L. Furia, and A. Diaspro, “Light-sheet confined super-resolution using two-photon photoactivation,” PLoS One 8, e67667 (2013).

28. L. Gao, L. Shao, B.-C. Chen, and E. Betzig, “3D live fluorescence imaging of cellular dynamics using Bessel beam plane illumination microscopy,” Nature Protocols 9, 1083–1101 (2014).

29. A. Escobet-Montalban, F. M. Gasparoli, J. Nylk, P. Liu, Z. Yang, and K. Dholakia, “Three-photon light-sheet fluorescence microscopy,” Opt Lett 43, 5484–5487 (2018).

30. G. Malkinson, P. Mahou, É. Chaudan, T. Gacoin, A. Y. Sonay, P. Pantazis, E. Beaurepaire, and W. Supatto, “Fast In Vivo Imaging of SHG Nanoprobes with Multiphoton Light-Sheet Microscopy,” ACS Photonics 7, 1036–1049 (2020).

31. M. Drobizhev, S. Tillo, N. S. Makarov, T. E. Hughes, and A. Rebane, “Absolute two-photon absorption spectra and two-photon brightness of orange and red fluorescent proteins,” J Phys Chem B 113, 855–859 (2009).

32. R. Kopp, T. Schwerte, and B. Pelster, “Cardiac performance in the zebrafish breakdance mutant,” The Journal of experimental biology 208, 2123–2134 (2005).

33. J. Gierten, C. Pylatiuk, O. T. Hammouda, C. Schock, J. Stegmaier, J. Wittbrodt, J. Gehrig, and F. Loosli, “Automated high-throughput heartbeat quantification in medaka and zebrafish embryos under physiological conditions,” Scientific Reports 10, 2046 (2020).

34. P. Mahou, J. Vermot, E. Beaurepaire, and W. Supatto, “Multicolor two-photon light-sheet microscopy,” Nat. Methods 11, 600–601 (2014).

35. M. Liebling, A. Forouhar, M. Gharib, S. Fraser, and M. Dickinson, “Four-dimensional cardiac imaging in living embryos via postacquisition synchronization of nongated slice sequences,” Journal of Biomedical Optics 10, 054001 (2005).

36. R. Amor, A. McDonald, J. Tragardh, G. Robb, L. Wilson, N. Z. Abdul Rahman, J. Dempster, W. B. Amos, T. J. Bushell, and G. McConnell, “Widefield Two-Photon Excitation without Scanning: Live Cell Microscopy with High Time Resolution and Low Photo-Bleaching,” PLoS One 11, e0147115 (2016).

