## Supplementary Material for "Balancing signal and photoperturbation in multiphoton light-sheet microscopy by optimizing laser pulse frequency"

### 1. Supplementary Methods

#### *1.1 Multiphoton light-sheet microscopy*

Multiphoton light-sheet microscopy (2P-SPIM) was performed using a custom-built optical set-up (Fig. 2a). The illumination arm was composed of an Ytterbium fiber femtosecond laser (Satsuma, Amplitude) producing pulse trains at 1030 nm wavelength, with 290-310 fs pulse duration and 0.6-40 MHz tunable frequency. Fast control of the illumination power was obtained with either an electro-optic modulator (EOM, Conoptics) or the laser built-in acousto-optic modulator, with additional wave plates and polarization beam splitters to tune the power modulation range. A shutter (Thorlabs) was used to completely block the laser. The illumination beam propagating in the x-direction (Fig. 2a) was focused on the sample by a low numerical aperture (NA) water immersion objective (10× 0.30 NA, Nikon). Illumination beam divergence was controlled with a telescope so as to achieve an effective illumination NA of ~0.09, corresponding to a Gaussian beam waist $w_0$ of ~3.6 µm (light-sheet thickness) and an axial resolution of ~3 µm of two-photon excited fluorescence signal. A pair of galvanometer mirrors produces on the one hand the illumination light sheet by vertical scanning of the beam in the y-direction synchronously with the camera read-out, and on the other hand the displacement in z of the light-sheet to perform volume imaging. The white light illumination was performed with a LED desk lamp focused onto the sample with a 10× 0.30 NA objective (Nikon) from the opposite side to that used for laser illumination. The detection arm was composed of a high NA water immersion objective (16× 0.80 NA, Nikon) collecting fluorescence orthogonally to the light-sheet plane. The z-position of the detection objective was moved in synchrony with the z-scanning of the light-sheet by using a piezo stage. A short-pass 800 nm filter was used to detect signal only in the visible range. A tube lens was used to image the illuminated plane onto a sCMOS camera (Flash 4.0, Hamamatsu) with a pixel size of 0.4 µm. The camera line detection was synchronized with the illumination y-scanning. The Sample was maintained in chamber filled with water solution and positioned from the top of the chamber with a combination of a motorized stage (MP285, Sutter Instrument) for translation in X, Y and Z directions, and a rotation stage for rotation about the Y axis. The sample was mounted in agarose using a glass capillary tube. Its spatial position was controlled by the combination of a rotation stage about the y-axis and a three-way motorized stage (MP285, Sutter Instrument), positioned over a water-filled chamber. All peripheral devices, including laser illumination power using the EOM, shutter, galvanometer mirrors, piezo-stage, sCMOS camera, and motorized sample stage, were controlled and synchronized using custom-written

LabVIEW (National Instruments) software. Image acquisition was performed at maximum imaging speed. It was determined by the vertical size (y-direction) of the field of view, with a camera line detection set to its minimum (9.8 µs), corresponding to the imaging speed of 168 frames per second for a 500 pixels field of view in the y-direction. In this case, each image was illuminated and detected in 4.9 ms, with 1.1 ms additional delay between images, during which the laser was switched off. The camera detection time per pixel was set to 490 µs. As a consequence, the signal per pixel was limited by the pixel illumination time during a y-scan, corresponding to ~73 µs/pixel. $P_{mean}$ corresponds to the mean power at the sample during imaging taking into account the laser switching off between successive images. To estimate $S_L$ depending on the illumination wavelength (Fig. 3c), a different home-built multiphoton light-sheet microscope was used as previously described in [34].

#### *1.2 Sample preparation for imaging*

The following zebrafish lines were provided by AMAGEN zebrafish facility at CNRS Gif-sur-Yvette (UMS 3504 CNRS / UMS 1374 INRA): *casper*, *casper* crossed with Tg(*ubi*:histone-mCherry) and *casper* crossed Tg(*ubi*:nls-TagRFP). Embryos were either obtained from AMAGEN or from the Ecole polytechnique zebrafish facility. Eggs were raised in the dark at 28°C for four to five days. For imaging, embryos were anaesthetized with 0.01% (100 mg/L) Tricaine (MS-222, Sigma Aldrich) and embedded in 1% (10 g/L) low melting point agarose (Sigma Aldrich) as previously described [4]. Imaging was performed at room temperature (20-23°C) in the imaging chamber filled with 0.01% Tricaine solution. All experiments were performed with zebrafish embryos before independent feeding larval forms and complied with the European directive 2010/63/UE. KTP nanocrystals were prepared and mounted for imaging as previously described [30]. Blue FluoSpheres (350/440, Thermofisher F8815) were diluted at a ratio of 1:1000 in agarose.

#### *1.3 Image analysis and reconstruction*

Image analyses were performed using Matlab (The MathWorks Inc.). To estimate the signal level (Fig. 1), cell nuclei or point sources were segmented using an intensity threshold and the signal averaged from all segmented pixels. 4D reconstruction (Fig. 6b and visualization 1) using post-acquisition time synchronization of beating heart images was performed as previously described [34, 35]. 3-D rendering and manual heart segmentation were performed using Imaris (Bitplane). For display of heart images (Fig. 6b and visualization 1), the pixel histogram was first linearly stretched and then gamma adjusted with a gamma value of 2.

#### *1.4 Heart beat rate analysis and definition of $S_L$ and $P_{NL}$*

To estimate $S_L$ and $P_{NL}$, the heart beat rate (HBR) was quantified depending on the illumination parameters. In each experiment described in Fig. 2, the heart was imaged at 168 fps using white light illumination and time series of 22 000 images (500x500 binned by a factor 20). The first 3000 images (18 s of acquisition) were used to quantify the HBR baseline ($HBR_0$). Then, during 10000 images (from acquisition time 18 to 77 s), the femtosecond laser was switched on at a given mean power $P_{mean}$ and period $T$ and the embryo was illuminated in the same manner as during 2P-SPIM imaging. The ΔHBR variation was quantified during this time. Finally, during the last 12000 frames (from acquisition time 77 to 131 s), the femtosecond laser was switched off to follow the HBR returning to the baseline in case the HBR variation was reversible. The instantaneous HBR was estimated using custom-made scripts written in Matlab (The MathWorks Inc.) as follows. A 10 second time-windowed fast Fourier transform (FFT) of the temporal signal from each pixel was performed (Fig. 2b-c). To obtain the instantaneous HBR, the 30 best pixels were selected based on their FFT signal-to-noise measure (SNR), defined as the ratio of the FFT maximum over the time window divided by the mean value of the FFT. Instantaneous HBR value at each pixel was estimated with sub-timepoint accuracy using a three-point interpolation (Fig. 2c). The mean HBR value from the 30 selected pixels at each time point was then estimated and plotted depending on time as in Fig. 2d. ΔHBR relative variation was defined as $HBR-HBR_0$, the difference between the HBR after and before switching the laser on. Finally, this experiment was repeated on the same embryo at a given laser pulse frequency *f* using an increasing $P_{mean}$. The reversible nature of the laser induced effect were verified by observing the HBR returning to the same $HBR_0$ at the beginning of each measurement (Visualization 1). $P_{mean}$ was increased up to the observation of irreversible photodamage such as heart arrhythmia or bubble formation (Visualization 1). The threshold $P_{NL}$ was then defined as the mean between the highest $P_{mean}$ used without irreversible photodamage and the $P_{mean}$ inducing them. Finally, $S_L$ was defined as the slope of the linear regression of ΔHBR depending on $P_{mean}$ at a given laser period $T$ (Fig. 2e). Such measurements were also performed depending on the illumination beam x-position within the heart (Fig. 3b and 4c) and on the illumination wavelength (Fig. 3c).

#### *1.5 Photobleaching analysis*

To quantify photobleaching depending on the laser period $T$, zebrafish embryo expressing mCherry were imaged in the tail region to avoid artefact due to cell motion. A different area was imaged in each condition. The photobleaching experiments were performed at constant fluorescent signal levels by adjusting $P_{mean}$ to keep $T.{P_{mean}}^2$ constant. Due to the Gaussian profile of the illumination in the x-direction, photobleaching is stronger at the center of the field of view. We then selected the center part of the image (40 µm large in the x-direction) to analyze the 2PEF signal decay. Finally, to optimize the exponential decay fit of the experimental data using scripts written in Matlab (The MathWorks Inc.), we analyzed 1000 images for *f*=0.6-2 MHz (fast decay), and 5000 images for *f*=5-40 MHz (slow decay). The photobleaching rate *k* is then defined as the decay rate of an exponential fit:

$$2PEF\ signal\ (image\ number) \sim \mathrm{A} + \mathrm{B} * \ e^{-k.image\ number}$$

#### *1.6 Signal enhancement graph*

To balance signal and photoperturbation, we plotted on the single graph how the 2PEF and 3PEF signal enhancement (Fig. 6a and Fig. S3, respectively) is limited by unwanted processes (heating, highly nonlinear photodamage and photobleaching). In such a graph, the $n_S$-order signal enhancement $S_{n_S}$ is plotted in logarithmic scale depending on the laser pulse period *T*. We then have:

$$S_{n_S}(T, P_{mean}) = \frac{T^{n_S-1}{P_{mean}}^{n_S}}{{T_0}^{n_S-1}{P_0}^{n_S}}$$

with $T_0$ = 1/80 MHz and $P_0$ = 70 mW, corresponding to typical laser parameters currently used in multiphoton light-sheet microscopes. In this graph, a line of constant $P_{mean}$ has a slope of $n_S - 1$. In the case of 2PEF or 3PEF signals, lines of constant $P_{mean}$ have a slope of 1 ($n_S = 2$, blue-to-red color dashed lines in Fig. 6a) or 2 ( $n_S = 3$, blue-to-red color dashed lines in Fig. S3).

In the case of a $n_P$-order photoperturbation, with a power threshold $P_{th}$(T), we have:

$$P_{th}(T) = \left(\frac{T}{T_0}\right)^{\frac{1-n_P}{n_P}} P_{th}(T_0)$$

and

$$S_{n_S}(T, P_{th}) = \left(\frac{T}{T_0}\right)^{\frac{n_S}{n_P}-1} \left(\frac{P_{th}(T_0)}{P_0}\right)^{n_S}$$

As a consequence, a line of constant photoperturbation has a slope of $\frac{n_S}{n_P} - 1$ in this graph. The sign of this slope is critical to balance the signal and the photoperturbation and depends on their relative order.

In the case of 2PEF signal ($n_S = 2$, Fig. 6a), a line of constant $n_p$-order photoperturbation has a slope of $\frac{2}{n_P} - 1$. This slope is positive and equal to one in the case of a linear effect such as heating ($n_P = 1$). Hence, lines of constant linear effect (blue-to-red color dashed lines in Fig. 6a) correspond to lines of constant $P_{mean}$ and to specific linear variation of HBR and of temperature, as estimated from our experimental investigation. In the case of $n_P > 2$, such as for photobleaching or nonlinear photodamage, the slope is negative (solid and dashed black lines in Fig. 6a, for $P_{NL}$, and photobleaching threshold, respectively). For instance, the nonlinear photodamage threshold $P_{NL}$ ($n_P$=5.8) reported in section 2.4 follows a line:

$$S_2(T, P_{NL}) = \left(\frac{T}{T_0}\right)^{\frac{2}{5.8}-1} \left(\frac{P_{NL}(T_0)}{P_0}\right)^{5.8},$$

with $P_{NL}(T_0)$~1100 mW is estimated from our experimental fit.

It is important to note that all these unwanted processes have an order that is different from *2*. As a consequence, adjusting the laser parameters has differential impact on signal of order 2 and on unwanted effects. In general, a reduction of photoperturbation of order $n_P < 2$, (respectively $n_P > 2$), is obtained by increasing (respectively decreasing) $T = 1/f$. In our graph, this is highlighted by positive (respectively negative) slopes of lines of constant photoperturbation. For instance, since highly nonlinear photoperturbations ($n_P > 2$) were often reported as the limiting factor in point-scanning multiphoton microscopy [16, 17, 22], it was proposed to increase *f* above 80 MHz to mitigate them [16]. Here, we observed the opposite behavior during live 2P-SPIM imaging. Indeed, the first observable effect is linear and is likely due to heating by water absorption, which calls for a different optimization approach.

### 2. Supplementary Figures

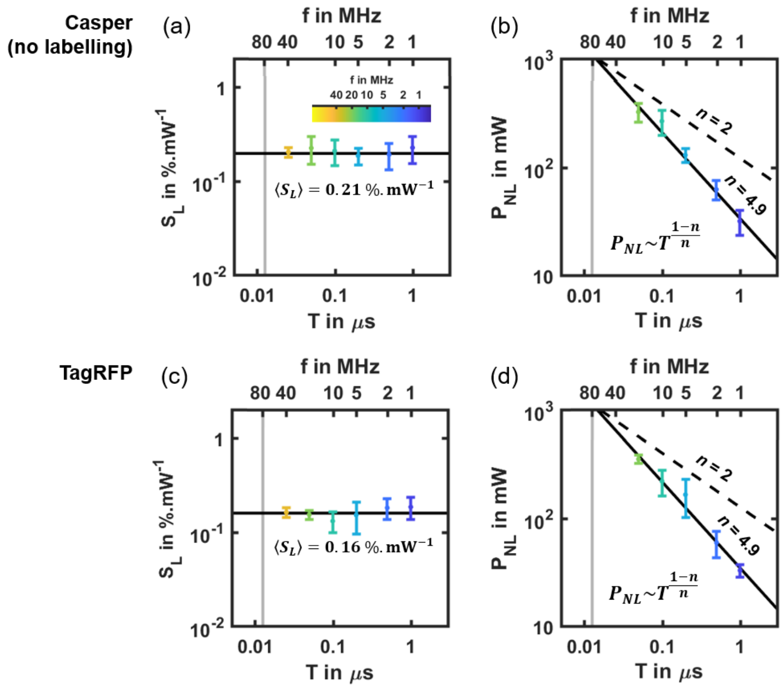


**Fig. S1.** Linear effect and nonlinear photodamage threshold in unlabeled and TagRFP labeled embryos. (a) Linear slope $S_L$ of the HBR relative variation of unlabeled *casper* zebrafish hearts (N = 32 embryos) depending on the laser pulse frequency $f = 1/T$. Black line indicates mean value $<S_L> = 0.211 \pm 0.017$ %.mW$^{-1}$). (b) Nonlinear photodamage threshold $P_{NL}$ in unlabeled hearts (N= 27 embryos) depending on the laser pulse frequency $f$=1/$T$. Black line shows the result of the scaling law fitted on logarithmic scaled data. The $P_{NL}(T)$ follows a scaling law of order $n$~4.9. (c) Linear slope $S_L$ of the HBR relative variation of TagRFP labeled zebrafish hearts (N = 27 embryos) depending on the laser pulse frequency $f = 1/T$. Black line indicates mean value $<S_L> = 0.161 \pm 0.014$ %.mW$^{-1}$). (d) Nonlinear photodamage threshold $P_{NL}$ in TagRFP labeled hearts (N = 23 embryos) depending on the laser pulse frequency $f = 1/T$. Black line shows the result of the scaling law fitted on logarithmic scaled data. The $P_{NL}(T)$ follows a scaling law of order $n$~4.9. Error bars indicate standard deviation. Black dashed line indicates a scaling low of order $n = 2$ to show how it deviates from 2PEF signal. Results of scaling law fits are listed in Table S1.

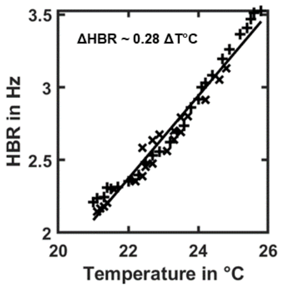


**Fig. S2.** Heart beat rate depending on temperature for 4-5dpf zebrafish embryos. The embryos were placed in warm water and their HBR was measured as the water cooled down. HBR increases linearly with temperature. Black line indicates the result of a linear fit (ΔHBR in Hz = 0.285 ± 0.015 ΔTemperature in °C, $R^2 = 0.97$).

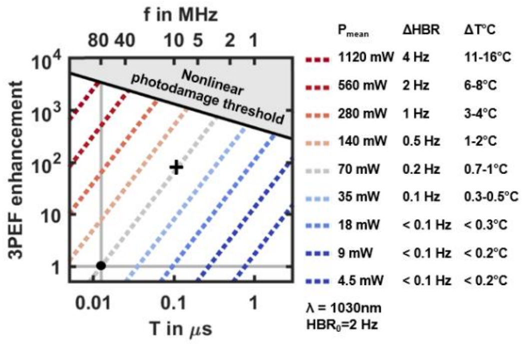


**Fig. S3.** 3PEF signal enhancement graph used to select the optimal laser pulse frequency. Signal enhancement corresponds to $\frac{T^2 P_{mean}{}^3}{T_0{}^2 P_0{}^3}$ with $T_0$=1/$f_0$=1/80 MHz and $P_0$=70 mW (black dot). Solid black lines correspond to the nonlinear photodamage threshold. Dashed blue-to-red lines corresponds to line of constant mean power of slope 2 and constant linear effect (variation of heart beat rate ΔHBR and of temperature ΔT°C). Black cross, imaging conditions used in Fig. 6b.

### 3. Supplementary Tables

| Figure | Experiment | Laser repetition rate $f$ | Laser mean power $P_{mean}$ | Laser scan speed | Field of view | Frame rate |
|---|---|---|---|---|---|---|
| | | MHz | mW | µm.ms$^{-1}$ | pixel$^2$ | Frame.s$^{-1}$ |
| 1a | $2PEF$ signal | 4.4<br>$to$<br>40 | 100 | 40 | $500 \times 500$ | 168 |
| 1b | $SHG$ signal | 0.6<br>$to$<br>40 | 100 | 8 | $2048 \times 2048$ | 41 |
| 1c | $3PEF$ signal | 4<br>$to$<br>13 | 54 | 40 | $500 \times 500$ | 33 |
| 4a | | 1<br>2<br>5<br>10<br>20 | 29<br>51<br>126<br>201<br>322 | 40 | $500 \times 500$ | 168 |
| S1b | Nonlinear photodamage threshold | 1<br>2<br>5<br>10<br>20 | 32<br>63<br>131<br>269<br>327 | 40 | $500 \times 500$ | 168 |
| S1d | | 1<br>2<br>5<br>10<br>20 | 33<br>60<br>166<br>221<br>353 | 40 | $500 \times 500$ | 168 |
| 5a | Photobleaching experiment | 0.6<br>2<br>5<br>10<br>20<br>40 | 15<br>28<br>45<br>63<br>90<br>127 | 40 | $500 \times 500$ | 168 |
| 6b<br>Visualization 1 | 4D heart *in vivo* imaging | 10 | 70 | 40 | $500 \times 500$ | 168 |

**Table S1.** Experimental parameters.

| Fig. | Optical effect | Scaling law | $P_{mean}$ | Sample | $A$ | $B$ | $R^2$ | $n$ | $n$: 90% conf. interval |
|---|---|---|---|---|---|---|---|---|---|
| | | $Optical\ effect \sim T^?$ | | | Linear regression: $log(Optical\ effect) = A\ log(T) + B$ | | | | |
| 1a | $2PEF$ signal | $2PEF \sim T^{n-1}$<br>$n = A + 1$ | $P_{mean} = cst$ | mCherry embryos | 1.2 | 2.1 | 0.91 | 2.2 | [1.9, 2.5] |
| 1b | $SHG$ signal | $SHG \sim T^{n-1}$<br>$n = A + 1$ | $P_{mean} = cst$ | KTP nanocrystals | 1.0 | 1.9 | 0.998 | 2.0 | [2.0, 2.1] |
| 1c | $3PEF$ signal | $3PEF \sim T^{n-1}$<br>$n = A + 1$ | $P_{mean} = cst$ | Fluo-spheres | 2.0 | 2.0 | 0.999 | 3.0 | [3.0, 3.1] |
| 3a | | | | mCherry embryos | 0.032 | 1.3 | 0.016 | 1.0 | [0.95,1.1] |
| S1a | Linear effect $S_L$ | $S_L \sim T^{n-1}$<br>$n = A + 1$ | $P_{mean}$ varies to estimate $S_L$ | Unlabeled embryos | $-0.009$ | 1.3 | 0.002 | 0.99 | [0.92,1.1] |
| S1c | | | | TagRFP embryos | 0.047 | 1.2 | 0.05 | 1.0 | [0.98,1.1] |
| 4a | | | | mCherry embryos | $-0.83$ | 1.5 | 0.98 | 5.8 | [4.4, 8.2] |
| S1b | Nonlinear photodamage threshold $P_{NL}$ | $P_{NL} \sim T^{\frac{1-n}{n}}$<br>$n = 1/(A+1)$ | $P_{mean} = P_{NL}$ | Unlabeled embryos | $-0.80$ | 1.5 | 0.93 | 4.9 | [3.6, 7.8] |
| S1d | | | | TagRFP embryos | $-0.80$ | 1.5 | 0.90 | 4.9 | [3.3, 9.8] |
| 5b | Photobleaching rate $r$ | $r \sim T^{\frac{n}{2}-1}$<br>$n = 2A + 2$ | $P_{mean} \sim T^{-2}$ | mCherry embryos | 0.67 | $-2.46$ | 0.99 | 3.3 | [3.2, 3.5] |

**Table S2.** Scaling laws of optical effects and estimation of their *n*-order using linear regression of logarithmic scaled data.

### 4. Visualizations

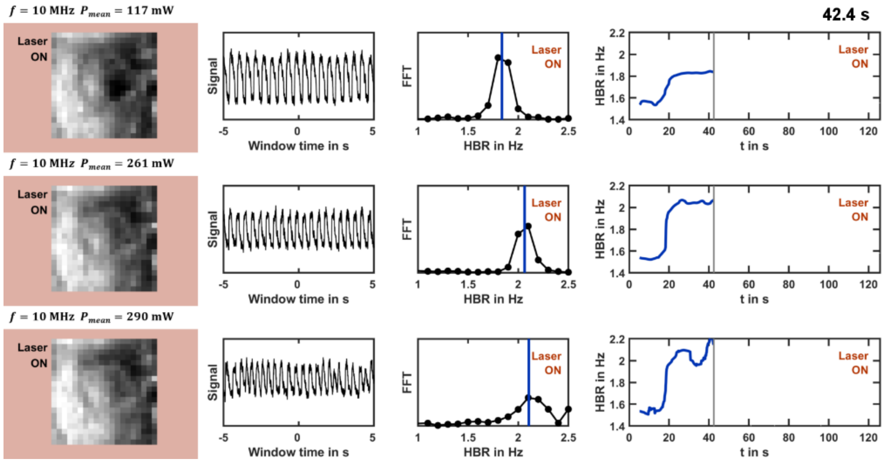


**Visualization 1.** Experimental workflow of HBR analysis. Presenting the workflow for three acquisitions, at $P_{mean}$ = 117 mW (top), 261 mW (middle) and 290 mW (bottom) on the same zebrafish heart at $f$ = 10MHz. First column: white light illumination images of the heart of the embryo. Second column: periodic signal fluctuation extracted from individual pixels over a 10 s window. Third column: windowed Fourier transform of the signal to extract of the HBR over that window. Bottom line: HBR as a function of time. Nonlinear photodamage are observed at $P_{mean}$ = 290 mW with heart beat arrhythmia followed by intense signals.

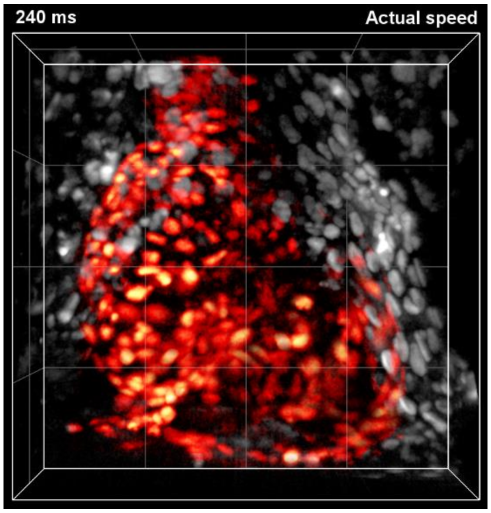


**Visualization 2.** 4D reconstruction of the zebrafish beating heart imaged with 2P-SPIM at 168 fps with optimized acquisition parameters. Histone mCherry-labeled zebrafish embryo imaged using $f$=10MHz and $P_{mean}$=70mW with 200x200x100 µm or 500x500x100 voxels field-of-view. Heart cells in red were manually segmented. Grid spacing of 50µm.
